# A SUMO interacting motif in the Replication initiator protein of Tomato yellow leaf curl virus is required for viral replication

**DOI:** 10.1101/2024.06.13.598784

**Authors:** Nicolas Frédéric Gaertner, Francesca Maio, Manuel Arroyo-Mateos, Ana P. Luna, Blanca Sabarit, Mark Kwaaitaal, Sandra Eltschkner, Marcel Prins, Eduardo R. Bejarano, Harrold A. van den Burg

## Abstract

CRESS-DNA viruses form a diverse group of viruses that use rolling-circle replication to replicate their genomes. They infect organisms in almost all branches of the eukaryotic tree of life. All CRESS-DNA viruses have one protein in common, the Replication initiator protein (Rep), which orchestrates viral replication using the host DNA replication machinery. In the case of the plant-infecting *Geminiviridae*, this multifunctional protein both recruits the host DNA replication machinery and manipulates posttranslational modification including Small ubiquitin-like modifier (SUMO) conjugation. In fact, Rep from two different geminiviruses, Tomato yellow leaf curl virus (TYLCV) and Tomato golden mosaic virus (TGMV), was shown to interact with the SUMO conjugating enzyme SCE1. Here, we demonstrate that also TYLCV Rep interacts with Arabidopsis SUMO1 and report on a SUMO interacting motif (SIM) in the SF3 helicase domain of Rep. Remarkably, an intact SIM proved to be important for the interaction of Rep with both SUMO1 and SCE1. The same motif was also essential for viral replication and Rep ATPase activity. Our findings thus connect the interaction between Rep and the SUMO machinery with viral replication of TYLCV.

**Importance:** The identification of a non-canonical SUMO-interacting motif (SIM) within the Rep protein of Tomato yellow leaf curl virus (TYLCV) reveals a connection between viral replication and a protein modification, SUMOylation. Importantly, the motif was found to be conserved between Rep proteins from different geminiviruses. Functionally, the motif was critical for the interaction of Rep with proteins of the SUMO machinery, viral DNA replication, and Rep ATPase acitvity. In particular, the third position of the motif was important for each of these activities. We thus uncover a novel mechanism on how geminiviruses recruit the SUMO machinery likely to their own need.

## Introduction

Circular replication (Rep)-encoding single stranded (CRESS)-DNA viruses form a unique group of viruses that use rolling-circle replication (RCR) to replicate their genomes (1–3). They have been found in almost all branches of the eukaryotic tree of life. The *Geminiviridae* family of viruses represents an important group of plant-infecting viruses that cause severe crop damages (4). Interestingly, CRESS-DNA viruses have only one protein in common, i.e. Replication initiator protein (Rep) that orchestrates RCR (5, 6). Thereto, these viruses need to recruit host factors including DNA polymerases to form viral replisomes with a largely mysterious but critical role for Rep (7–11). At the same time, Rep manipulates cellular processes by steering protein modifications like Ubiquitin or Small ubiquitin-like modifier (SUMO) conjugation (SUMOylation) onto host proteins (12–14). By now it is well established that many viruses manipulate host SUMOylation to suppress anti-viral responses or to enhance viral replication (15). Yet, how and why Rep recruits and directs SUMOylation via one or more target proteins remains enigmatic.

Three protein domains can be identified in Rep. Its N-terminal domain, called the HUH (His-hydrophobic residue-His) endonuclease domain, nicks and joins the single-stranded viral genome at the origin-of-replication using an active-site tyrosine, that forms a 5′-phosphotyrosine bond with the nicked DNA (16). The central domain is involved in Rep oligomerization (17–19), whereas the C-terminal domain shares homology with the ATPase helicase superfamily 3 (SF3 helicase) and is composed of a canonical Walker A and B motif. This domain is presumed to act as a replicative helicase during RCR elongation (19–24). It was reported earlier that Rep from different geminiviruses is also able to reprogram the host cell cycle thereby reactivating DNA replication in terminally differentiated plant cells (25, 26). Thereto, Rep interacts with a multitude of host proteins implicated in the cell cycle (notably DNA replication) and in DNA repair. For example, Rep was found to interact with transcriptional regulators like Retinoblastoma-related protein (RBR) (27, 28), but also with Proliferating cell nuclear antigen (PCNA), Replication factor C (RFC), Replication protein A32 (RPA32), and Minichromosome maintenance protein 2 (MCM2) (29–32).

The general notion is that by the process of SUMOylation entire protein complexes become decorated with SUMO at multiple sites, rather than modifying a single site of one particular protein substrate (33). In this way, SUMOylation modulates a wide range of nuclear processes including the cell cycle, DNA replication, DNA damage repair, RNA processing, and gene expression (34). SUMO attachment involves a cascade of enzymatic reactions that starts with (i) precursor maturation, followed by (ii) SUMO activation by the SUMO E1-activating enzyme (SAE1/SAE2 dimer), and (iii) SUMO transfer to target proteins by the SUMO E2-conjugating enzyme (SCE1) (35–39).

Importantly, in the case of two geminiviruses, i.e. Tomato yellow leaf curl virus (TYLCV) and Tomato golden mosaic virus (TGMV), Rep was confirmed to interact with the SCE1 protein (12, 14, 40). Furthermore, in the presence of Rep, SUMOylation of PCNA was impaired, both *in vitro* and *in vivo* (14). At least in yeast, loss of PCNA SUMOylation is known to cause an increase in homologous recombination (41), a process critical for geminivirus DNA replication (42). Two Lys residues in the HUH endonuclease domain of Rep^TGMV^ proved to be essential for SCE1 binding and viral replication/spread inside the host plant (40). However, the same lysines were not essential for the interaction between Rep^TYLCV^ and SCE1 (43). Notwithstanding, Rep^TYLCV^ was found to interact with SCE1 in the nucleus where it formed protein aggregates called nuclear bodies (NBs) (43).

Here, we report how Rep^TYLCV^ interacts with Arabidopsis SUMO1 by revealing the existence of a previously unknown non-canonical SUMO-interacting motif (SIM) in the SF3 helicase domain. Our data imply that this SIM allows for the interaction with both SUMO1 and SCE1. We show that viral replication is compromised when this non-canonical motif is mutated. As this SIM is positioned next to the Walker A motif, we additionally assessed if mutating this motif would suppress Rep ATPase activity. We thus demonstrate that the SIM not only promotes the interaction with proteins of the SUMO pathway, but that it is also critical for ATPase activity and viral replication.

## Results

### Rep from TYLCV interacts with SUMO1 and SCE1 via a non-canonical SIM in the SF3 helicase domain

Previously, two Rep proteins (from TGMV and TYLCV) were found to interact with SCE1 (12, 14, 40, 43). As proteins involved in SUMOylation often interact with SUMO as well, we hypothesized whether (i) TYLCV Rep (hereafter just ‘Rep’, unless mentioned otherwise) also interacts with SUMO, (ii) if this interaction involves an unknown SIM in Rep, and (iii) if this is the case, whether this interaction somehow promotes the interaction with SCE1. To test this, the split-ubiquitin Y2H system (SUS Y2H) was used rather than the more frequently used GAL4 Y2H system (**Fig. 1A**), as the latter displayed autoactivation when Arabidopsis SUMO1 was expressed as bait protein (BD-SUMO). We found Rep to interact with both the mature protein (GG) and a conjugation-deficient variant (ΔGG) i.e., a variant that lacks the C-terminal diGly motif required for SUMOylation. This implies that a hitherto unknown SIM is present in Rep. To further explore this notion, we examined if Rep can still interact with a SUMO1 variant in which two residues critical for binding the SIM peptide are mutated into Ala (F32/I34A) (**Fig. 1B**) (63). Indeed, this SUMO1 variant (F32/I34A) failed to interact with Rep (**Fig. 1A**).

**FIG 1.**
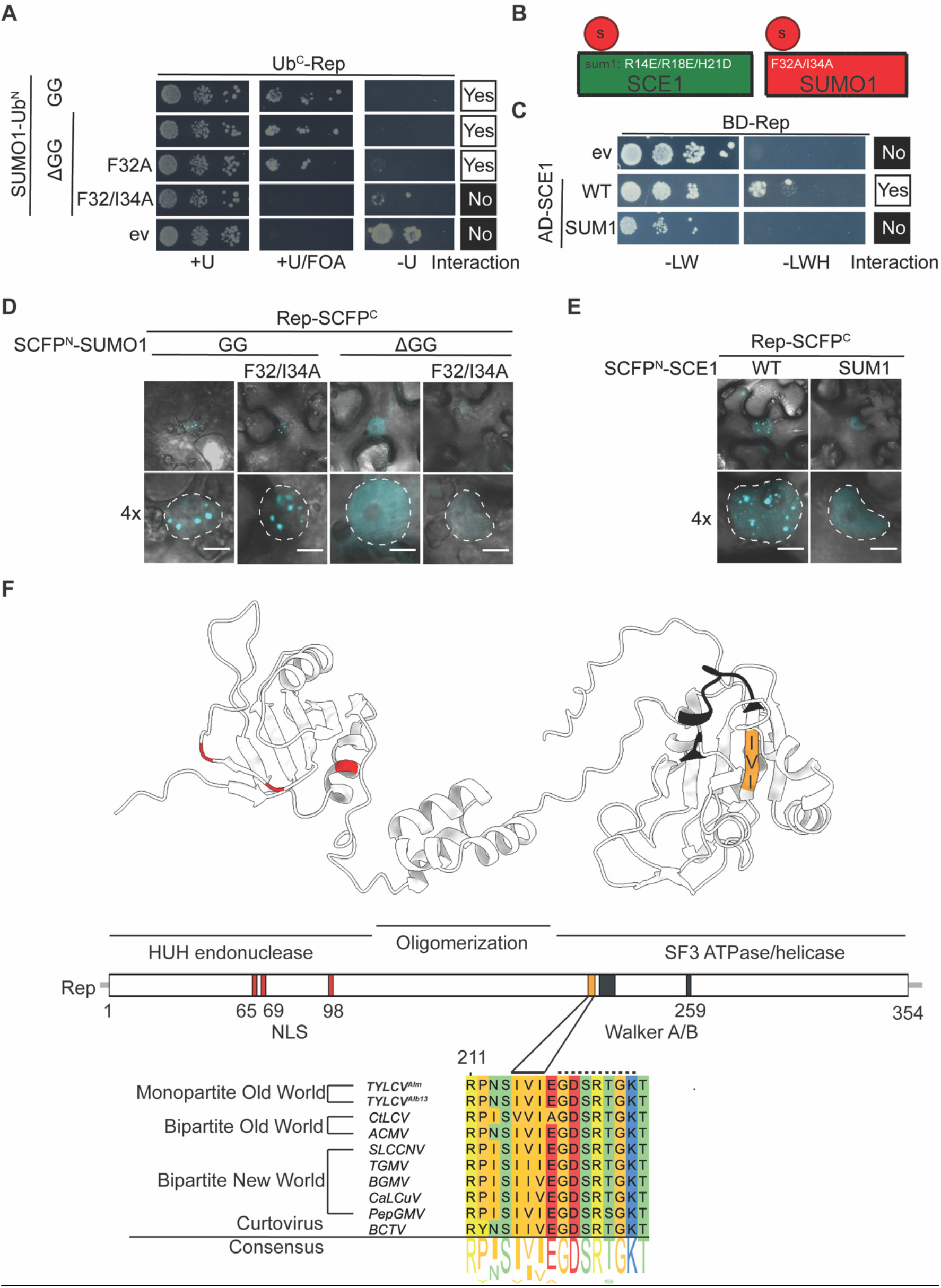
Rep interacts with SUMO1 and SCE1 possibly via a non-canonical SUMO interacting motif (SIM). (**A**) Yeast split-ubiquitin interaction assay between wildtype Rep (fused to C-terminal half of Ubiquitin, Ub^C^) and different SUMO1 variants (Ub^N^). Rep interacts with both the mature and a conjugation-deficient variant of SUMO1 (GG and ΔGG, respectively), but not when the SIM-binding pocket is mutated in SUMO1^ΔGG^ (F32A and F32/I34A). Positive interactions give growth on minimal medium (MM) supplemented with Uracil and FOA (+U/FOA) and no growth on MM lacking Uracil (−U). +U, positive control for normal growth. (**B**) Schematic representation of the SUMO1 and SCE1 variants used for Y2H and BiFC assays in panels A/D and C/E, respectively. (**C**) Gal4 yeast two-hybrid interaction between wildtype Rep_Alb13_ and two SCE1 variants: wildtype and variant mutated in the second SUMO binding site (sum1). Rep_Alb13_ interacts with wildtype (WT) SCE1 but not SCE1^sum1^. −LWH, growth when bait and prey interact; −LW, positive control for normal growth. (**D,E**) BiFC showing that Rep_Alb13_ interacts *in planta* with wildtype SUMO1 (D) and SCE1 (E) in NBs. Fluorescence reconstitution inside NBs is suppressed when Rep_Alb13_ is co-expressed with SUMO1^ΔGG+F32/I34A^ (panel **D**) or SCE1^sum1^ (Panel E). Images depict a typical *N. benthamiana* epidermal cell (top) and a 4× zoom showing only the nucleus (bottom). Dotted lines outline nuclei; scale bar, 5 μm. (**F**) Rep structure prediction using AlphaFold2 showing the functional domains. *Left-to-right*: HUH endonuclease, central oligomerization domain and Super Family 3 (SF3) ATPase/helicase domain. Residues critical for Rep nuclear localization (NLS), *red*; SIM, *orange*; Walker A and B motifs, *black*. Protein sequence alignment of Rep from different geminiviruses. Sequence shown comprises the residues 211-226. Residues 215-217 of Rep^Alm^ and 216-219 of Rep_Alb13_ represent the SIM (*continuous line*) and residues 219-225 depict the Walker A motif (*dashed line*).

Earlier it was shown that SCE1 can bind a second SUMO protein non-covalently via a site that is distal from the SCE1 catalytic pocket (63, 64). To assess if this second binding event is needed for Rep to interact with SCE1, a SCE1 variant (R14E/R18E/H20D/H21D; SCE1^sum1^) that has lost its ability to interact with SUMO1^ΔGG^ was used in the GAL4 system (**Fig. 1B**)(63). As hypothesized, we found that Rep binds the wildtype (WT) SCE1 protein, but not SCE1^sum1^ (**Fig. 1C**). These data suggest that at least in yeast the formation of a stable Rep-SCE1 complex depends (in part) on the interaction of SCE1 with SUMO1 as well. On this, we would like to note that yeast homologues of these proteins could possibly cross complement the plant homologs of these proteins – something that was not further investigated here.

Next, we investigated whether the SIM binding pocket of SUMO1 is also required *in planta* for the Rep-SUMO interaction. Using a Bimolecular fluorescence complementation (BiFC) assay we demonstrated earlier that (i) SUMO1 and SCE1 form large protein aggregates termed NBs and that (ii) NB formation was strictly dependent on recruitment of a functional SUMO conjugation enzyme to these bodies (63). Similar to the SUMO1-SCE1 interaction, we find that firstly Rep interacts with both SUMO1 and SCE1 inside NBs in the BiFC assay and that secondly this reconstitution of the fluorophore halves inside NBs was strongly suppressed in the case a fusion protein with SUMO1 ^ΔGG+F32/I34A^ (**Fig. 1D**) or SCE1^sum1^ (**Fig. 1E**). These data strongly suggest that Rep interacts with SUMO1 and possibly SCE1 via a SIM present in Rep.

Classically, SIMs are characterized by a stretch of at least three long-branched aliphatic residues, [VIL]-X-[VIL][VIL] (where ‘X’ denotes any residue), often directly flanked by a stretch of negatively charged residues (i.e. Glu, Asp, or phosphorylated residues) (44–48). To screen for SIMs in Rep, the tool GPS-SUMO was used (https://sumo.biocuckoo.cn). This yielded one non-canonical SIM motif in the SF3 helicase domain that was composed of a stretch of three long-branched aliphatic residues, i.e., Ile215, Val216 and Ile217. This putative SIM does not overlap with any other known viral open reading frame (ORF) of TYLCV (65). Yet, this SIM is located adjacent to the Walker A motif of the SF3 helicase domain (**Fig. 1F**). This means that this motif is in a different domain than the HUH endonuclease domain, which was already known to be important for the Rep-SCE1 interaction. A pan-genome comparison of the geminiviruses revealed that this aliphatic stretch is widely conserved in the coding sequence of Rep, including Rep from both mono- and bipartite begomoviruses. (**Fig. 1F**)

To assess if this SIM binding pocket is required for the interaction with SUMO1^ΔGG^ and SCE1, the two Y2H systems described above were used again. To systematically test both interactions, the three long-branched aliphatic residues were mutated to Ala, as single residue mutations and in all possible combinations. Using the SUS Y2H assay, we found that the Rep single and double mutants I215A, V216A, I215/V216A, and I215/217A were still capable of interacting with SUMO1^ΔGG^, while the variants I217A, V216/I217A and the triple mutant I215/V216/I217A (hereafter Rep^sim^) did not (**Fig. 2A**). In the case of the Rep-SCE1 interaction, the double mutant I215/V216A or all Rep mutants that included I217A mutations failed to interact with SCE1 (**Fig. 2B**). To confirm these findings *in planta,* we resorted to the BiFC assay. As indicated above, the formation of SUMO NBs is considered to be a proxy for presence of SUMO-conjugation enzyme activity in SUMO-containing NBs (**Fig. S1A** and **S1B**) (63). Confirming the findings of the Y2H screen, we found that the Rep^sim^ triple mutant failed to aggregate inside NBs when co-expressed with SUMO1 (**Fig. 2C** and **S1C**) or SCE1 (**Fig. 2D** and **S1D**). As an independent method to test the interaction *in planta*, the split luciferase assay was employed. It offers the advantages of increased sensitivity and improved quantification of the interaction (66). Co-expression of wildtype Rep with SUMO1^ΔGG^ or SCE1 resulted in reconstitution of the luciferase protein resulting in an enhanced chemiluminescence signal. In contrast, co-expression of Rep^sim^ +SUMO1^ΔGG^ (**Fig. 2E**) or Rep^sim^ +SCE1 (**Fig. 2F**) gave a low chemiluminescence signal close, or comparable to, the background. Combined, these data suggest that the SIM motif is important for the interaction of Rep with SUMO1 and SCE1 both in yeast and *in planta*.

**FIG 2.**
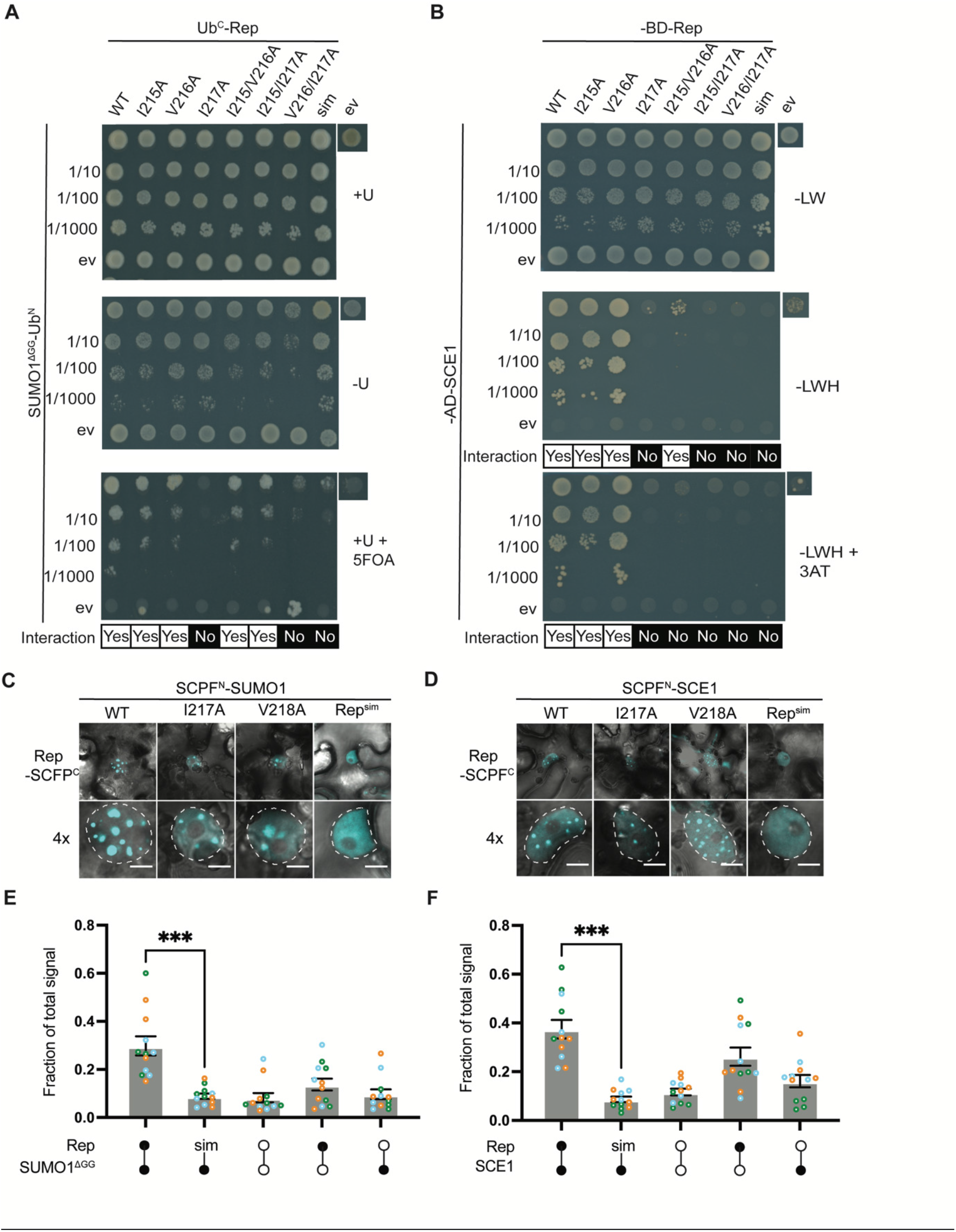
Rep SIM is needed for Rep to interact with both SUMO1 and SCE1. (**A**) Yeast split-ubiquitin interaction assay between Rep SIM mutants (Ub^C^) and SUMO1^ΔGG^(Ub^N^). Wildtype (WT) Rep and the variants I215A, V216A, I215/V216A and I215/217A interact with SUMO1^ΔGG^, while I217A, V216/I217A, and I215/V216/I217A (sim) do not. ev, empty vector control; *top-to-bottom*: 10-fold dilution series of the yeast cultures. Similar to the legend of Fig. 1A. (**B**) Gal4 yeast two-hybrid interaction between Rep SIM variants (GAL4 BD) and SCE1 (GAL4 AD). WT Rep and the variants I215A and V216A interact with SCE1, while RepI217A, the double and triple (sim) mutants do not. ev, empty vector control; top-to-bottom: 10-fold dilutions series of the yeast cultures. Similar to the legend of Fig. 1C. (**C,D**) BiFC showing SUMO1 (**C**) and SCE1 (**D**) localization pattern in complex with Rep_Alb13_ and the Rep _Alb13_ SIM variant. Rep^sim^ fails to form NBs in combination with SUMO1 (**C**) and SCE1 (**D**). Nuclei are outlined by a dashed white line; scale bars represent 5 μm. nls: nuclear localization-deficient variant (43). (**E**) Split-luciferase complementation assay between different Rep variants and SUMO1^ΔGG^. Only WT Rep in complex with SUMO1^ΔGG^ gave an enhanced chemiluminescence signal (open circles). Each dot in the bar graph represents a technical replicate, while colours represent independent biological repeats. X-axis: black dots represent expression of the indicated Rep/SCE1 variants. Open circles represent expression of empty vector controls for either half of the split-luciferase construct (Luc^C^/Luc^N^). ANOVA followed by a Dunnett multiple comparisons test was performed; *** p<0.001. Error bar represents standard error of the mean (SEM) (n=12). (**F**) Split-luciferase complementation assay between different Rep variants and SCE1. Only WT Rep in complex with SCE1 resulted in a chemiluminescence signal above the background (open circles). Other details as panel E.

### Rep SIM is essential for DNA replication activity in N. benthamiana

Next, we assessed if the Rep SIM was needed for viral DNA replication. Thereto, we first used a virus-free plant reporter system that mimics TYLCV replication (*2IR-GFP N. benthamiana*; **Fig. 3A**) (67). In this system, transient expression of Rep (in absence of other viral proteins) is sufficient to promote RCR of the *2IR-GFP* transgene cassette thereby forming circular ssDNA extrachromosomal molecules (ECMs) habouring a *GFP* expression cassette (*35Spro::GFP*) (67). With this system, we systematically tested whether introducing alanines in the SIM disrupted Rep DNA replication activity (**Fig. 3B**). To confirm both Rep protein stability and correct nuclear localization (43, 68) of the different variants, RFP was fused to the C-terminus (Rep-RFP). Mutating the SIM did not change the nuclear localization of the three Rep-RFP variants tested (**Fig. S1E**) (43,68). We also noted that upon transient expression *in planta* the different Rep-RFP proteins accumulated to varying but detactable levels (**Fig. 3C, 3E,** and **3H; Fig. S1F**). When examining DNA replication activity, it was noted that the Rep variants I215/217A, V216/I217A and I215/V216/I217A displayed no discernable DNA replication activity (**Fig. 3D**). Although the detection of RFP fluorescence denotes expression of these variants (**Fig. 3E** and **3H**), both the level of GFP fluorescence (**Fig. 3F**) and ECMs (**Fig. 3G**) were low compared to leaf discs expressing wildtype Rep (**Fig. 3D** and **3F**). In order to compare the relative GFP levels, the GFP signals were adjusted according to the Rep-RFP levels (**Fig. 3H**); please note that low *GFP* expression are already occurs in absence of Rep due to the 35S promoter. Based on these observations, it is evident that the SIM is needed for DNA replication activity, with Ile217 having the strongest contribution in combination with Ile215 or Val216.

**FIG 3.**
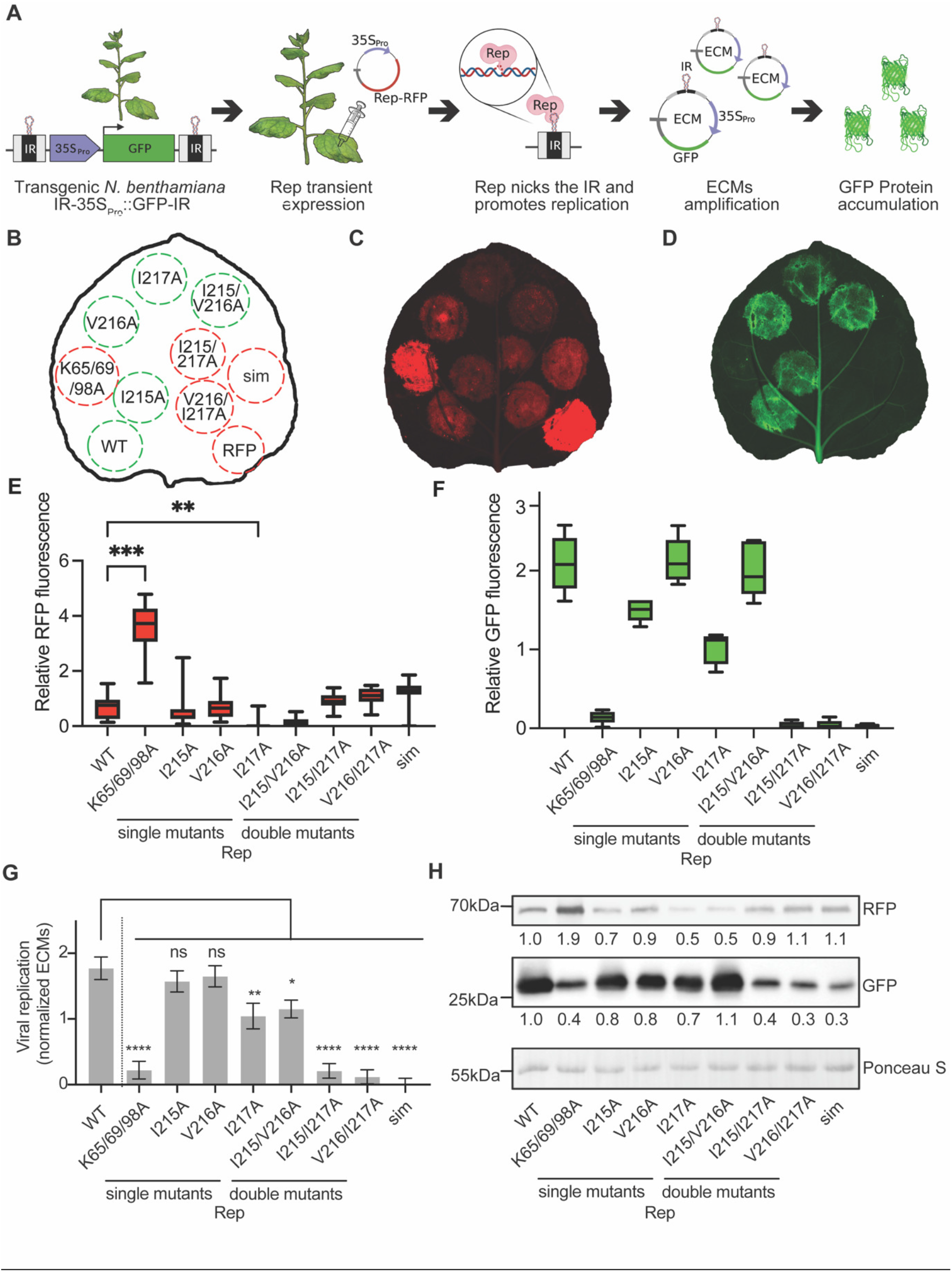
Rep SIM is important for viral DNA replication in a virus-free reporter system. (**A**) *N. benthamiana* plants expressing the *IR-35Spro::GFP-IR* reporter to visualize viral replication via increased GFP fluorescence. Transient expression of Rep causes DNA-nicking of the IR motif and promotes RCR of the nicked DNA resulting in formation of circular ECMs. Each ECM contains a *35Spro::GFP* cassette resulting in massive GFP accumulation. (**B**)Scheme of the infiltrated constructs; bottom/left clockwise: WT Rep; K65/69/98A (nls), replication-deficient negative control impaired in nuclear localization (43); different SIM mutants, single (I215A, …), double (I215/V216A, …), triple residue (sim) mutants, and RFP negative control. Green and red circles depict replication-competent and -deficient Rep variants, respectively. (**C**) Fluorescence image depicting Rep-RFP accumulation. Panel B depicts layout of the infiltrated constructs. (**D**) Fluorescence image depicting GFP accumulation as proxy for replication. Same leaf as panel C; (**E, F**) RFP (E) and GFP (F) Quantification of fluorescence signal in infiltraded leaves. Total fluorescence was normalized per leaf. Kruskal-Wallis test with Dunn’s correction was performed on the pooled data from each replicate. ** p<0.01; *** p<0.001. Error bar represents SEM (n=14). (**G**) ECM quantification using real-time PCR. ANOVA followed by Dunnett’s multiple comparison test between was performed; * p<0.05; ** p<0.01; *** p<0.001; **** p<0.0001. Error bar represents SEM (n=3). (**H**) Immunoblot showing both Rep-RFP and GFP levels. Shown are ratios compared to signal intensity of WT Rep. To demonstrate equal protein loading and normalize the blot signal values, membranes were stained with Ponceau S.

### Rep SIM is also required for TYLCV replication and viral spread in plants

Although Rep DNA-replication activity was strongly impaired by introducing alanines in the SIM, we wondered whether introduction of the same mutations in an infectious viral clone would also prevent viral replication. To this end, a TYLCV infectious clone was used as previously reported (69). Three mutant variants of this clone were generated that contained different Ala substitutions in the Rep SIM, while leaving all other known viral ORFs intact (65). Based on the residual REP DNA replication activity, we hypothesized that the TYLCV variant I215/V216A would still yield an active virus, while the mutations I215/217A and I215/V216/217A (TYLCV^sim^) should give replication-deficient clone (‘non-infectious’). Indeed, agroinfiltration of *2IR-GFP N. benthamiana* plants with wildtype TYLCV gave clear viral symptoms after three weeks including leaf curling, leaf puckering, and stunted plant growth (**Fig. 4A**). Also, TYLCV I215/V216A gave viral symptoms but to a lesser degree, which is in agreement with the lowered replication activity of Rep I215/V216A. TYLCV I215/217A and TYLCV^sim^ gave no viral symptoms analogous to loss of DNA replication activity by the corresponding Rep variants. Real-time PCR analysis confirmed these findings (**Fig. 4B**), i.e., the viral DNA titres were relatively low for TYLCV I215/I217A and TYLCV^sim^ in comparison to the wildtype virus. For TYLCV I215/V216A, the viral titres were one magnitude lower compared to wildtype TYLCV. These findings collectively demonstrate that Rep SIM is critical for efficient viral replication and spread, and that in particular residue Ile217 is critical.

**FIG 4.**
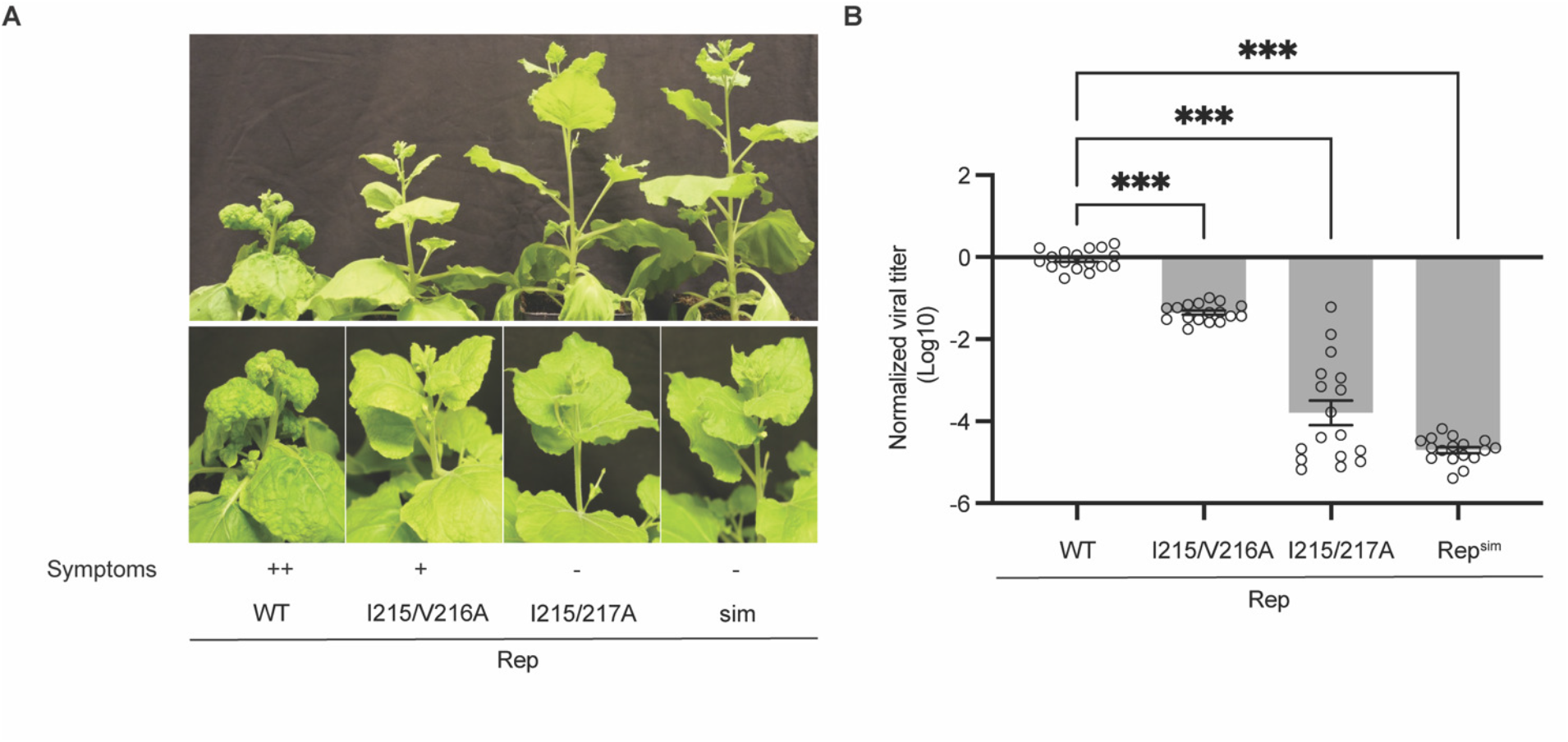
Also in the viral context the Rep SIM is required for viral replication and viral spread in *N. benthamiana.* (**A**)Picture of *2IR-GFP N. benthamiana* upon infection with different TYLCV infectious clones expressing WT Rep or Rep SIM variants (I215/V126A, I215/I127A, I215/V126A/ I127A=sim). WT Rep infectious clone infected plant shows severe viral symptoms (++, severe symptoms; +, mild symptoms; −, no symptoms). Photos were taken three weeks post inoculation. (**B**) Quantification of viral titres using real-time PCR on the extracted DNA from the apical leaves. Levels were normalized to the viral titres by the wildtype virus (Log_10_=0). ANOVA followed by Dunnett’s multiple comparison test was performed; *** p<0.0001. Error bar represents SEM (n=17).

### Mutating the SIM impairs Rep ATPase activity and oligomerization

Due to the close proximity to the Walker A motif, the mutations in SIM could also affect Rep ATP activity (20) (**Fig. 1G**). To investigate this, we expressed and purified several Rep variants and tested ATPase activity. Specifically, we focused on the I217A mutations, considering the involvement of this residue both the SUMO/SCE1 interaction and viral replication (**Fig. 2** and **4**). Wildtype Rep displayed a high relative ATPase activity compared to different SIM mutants tested. Interestingly, Rep K225A, known for its impaired ATPase activity (20), showed more ATPase activity than any of the SIM mutants tested (**Fig. 5A**), which could be due to co-purification of *E. coli* chaperones. Overal, these observations suggest that the I217A mutation impairs Rep ATPase activity. Earlier studies on Rep from Porcine circovirus 2 highlighted its oligomerization into a hexamer complex bound to DNA (21). Notably, in the reported 3D structure (PDB: 7LAR, 7LAS) the ATP binding pocket is situated between two Rep subunits. Based on this finding, we hypothesized that the different SIM mutations could reduce ATPase activity by attenuating the capacity of Rep to form oligomers. To test this, we examined whether wildtype Rep exhibited a similar migration behavior to the SIM variants when loading them on a Size Exclusion Chromatography (SEC) column (**Fig. 5B**) and analyzing the protein composition of the different elution fractions using tandem mass spectrometry (MS/MS). The MS/MS analysis suggested that Rep did not elute in the fraction around the 8 mL mark (white arrow, estimated mass of 1.3 MDa), i.e., not a single MS peptide match was obtained with Rep (**Fig. 5B** and **Fig. S2**). In constrast, Rep was found to reside in the fractions that eluted around 16 mL (black arrow), corresponding to an approximate mass of 40 kDa (**Fig. 5B** and **Fig. S2**). Intriguingly, wildtype Rep had a more prominent shoulder at 14 mL in the elution profile compared to the Rep I217A and Rep I215/I217A, indicative for the formation of higher order complex. This shoulder at an elution volume of 14 mL could potentially correspond to an oligomeric state of Rep. However, at this stage, no definitive correlation could be drawn between the SIM mutation and its impact on the oligomerization status of Rep.

**FIG 5.**
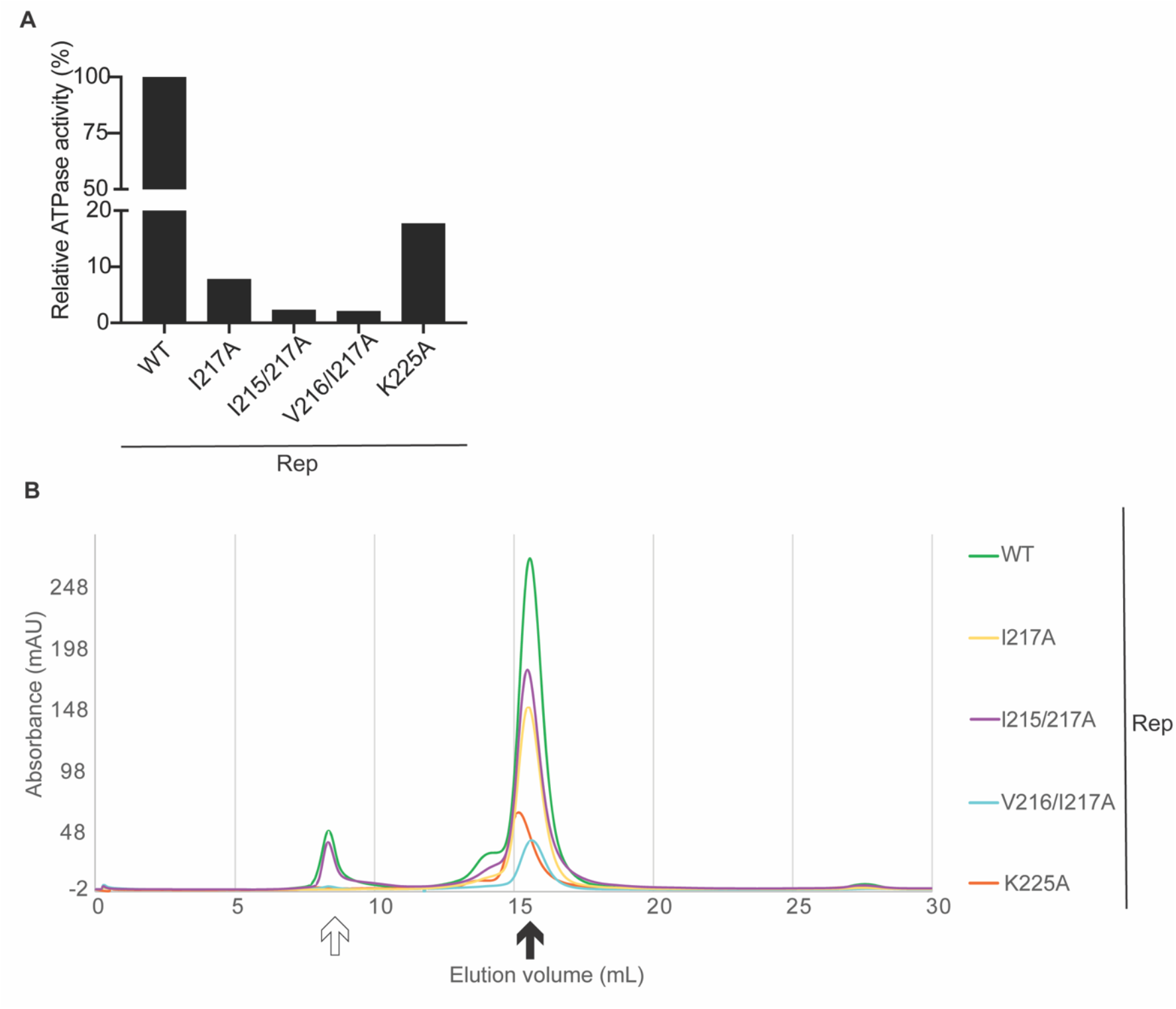
An intact SIM is needed for Rep ATPase activity. (**A**) Relative ATPase activity from pooled eluted fractions (approximate mass 40kDa) corresponding to wildtype (WT) Rep (100%) or Rep SIM mutants expressed as E. coli recombinant proteins. K225A: abolished ATPase activity Rep (negative control, ref. (20)). (**B**) Analytical size exclusion chromatography (SEC) of Rep SIM variants. The SIM variants display no clear difference in their oligomer state compared to WT Rep. Black and white arrow mark the protein fraction eluting at an apparent mass of 40 kD and 1.3 MDa, respectively.

## Discussion

CRESS-DNA viruses share one protein, Rep, that orchestrates their viral DNA replication. Previous studies reported that Rep from TGMV interacts with the SUMO E2 conjugating enzyme SCE1 via its N-terminal HUH endonuclease domain (12, 14, 40). Here we established that the SUMO protein itself is apparently involved in this interaction between Rep and SCE1. We provided evidence for the existence of a non-canonical SIM in the SF3 ATPase/helicase domain of Rep. This SIM was not only required for the interaction between Rep and Arabidopsis SUMO1, but also SCE1. Using a virus-free reporter system for viral replication (*2IR-GFP N. benthamiana*) (67), we demonstrated that replacing two residues in this SIM motif for alanines was sufficient to inhibit RCR by Rep, i.e. the third position in combination with the first or second position. Likewise, mutating this first and third position was sufficient to prevent viral replication of an infectious clone of TYLCV, demonstrating that none of the other viral transcripts or translated proteins was able to compensate for the introduced mutations in the Rep ORF. The relevance of this SIM is emphasized by the fact that it is conserved in Rep from different Geminiviruses, including both mono- and bipartite begomoviruses (**Fig. 1G**). This suggests that the residues of this motif have a conserved role in the viral life cycle.

The need for an intact SIM for the interaction with SUMO1/SCE1 was shown both in Y2H assays and *in planta* applying two independent assays (BiFC and the split-luciferase interaction). Earlier it was reported that overexpression of Rep did not result in a global change in the SUMO conjugate levels *in planta* (40). Here we make a related observation that, at least in the BiFC system, the protein pair Rep-SCE1 resides in NBs, similar to the SUMO1-SCE1 protein pair. As the formation of SUMO1-SCE1 NBs strictly depends on to presence of SCE1 enzymatic activity in these bodies (63), the formation of NBs corroborates again that Rep does not suppress SCE1 enzyme activity (**Fig. 2C** and **2D**).

In particular residue Ile217 appears to be pivotal for the interaction of Rep and SUMO1/SCE1. Most single and double mutants of the SIM were still capable of producing a protein that interacted with SUMO1^ΔGG^ except for Rep I217A, V216/I217A, and the triple mutant I215/V216/I217A (**Fig. 2A**). In the case of the interaction between Rep and SCE1, the double mutant I215/V21A and any mutant carrying the I217A subsitution was found to disrupt this interaction (**Fig. 2B**). The I217A mutation caused also a stronger reduction in viral replication (i.e. GFP expression from the ECMs and accumulation of viral DNA) than the Ile215 or Val216 single mutations (**Fig. 3F**). This establishes a novel function attributed to the SF3 ATPase/helicase domain, that is, SUMO binding with a critical role for the third position of the SIM.

The SF3 helicase domain comprises a Walker A and B motif required for Rep ATP binding and ATPase activity. When Lys225 of the SF3 ATPase/helicase domain is mutated into Ala, Rep ATPase activity was reported to be strongly impaired (20). The SIM identified here was also required for Rep ATPase activity. The latter would explain the loss of RCR activity by Rep, as ATPase activity is a prerequisite for Rep helicase activity (70). Yet, it is striking that this three-residue motif contributes to both the SUMO1/SCE1 interaction and viral replication. At this point we cannot exclude that suppression of PCNA SUMOylation by Rep involves this SIM (14). Additional research will be needed to elucidate the exact mechanisms through which the SIM impacts not only the interaction with SUMO1/SCE1, but also the ATPase activity of Rep.

Finally, we investigated the impact of mutating Ile217 on Rep oligomerization. Tarasova *et al*. (2021) (21) reported the presence of oligomers for Rep from Porcine circovirus 2. By size exclusion chromatography (SEC) analysis, a distinct shoulder was observed in the elution profile of wildtype Rep, that was absent for Rep I217A and I215/I217A (**Fig. 5B**). This shoulder likely corresponds to an oligomeric state of Rep. Yet, the expected hexameric conformation of approxmiately 324 kDa (**Fig. S2**), analogous to Porcine circovirus 2 Rep, could not be substantiated under the experimental conditions used. Therefore, no definitive correlation can be drawn between the SIM mutations and their impact on Rep oligomerization.

## Materials and methods

### General methods and cloning

All molecular techniques were performed using standard methods (49). *Escherichia coli* strain DH5α was used to clone gene fragments. All primers and plasmids used are described in the **Tables 1** and **2**, respectively. The different fragments were PCR-amplified using Phusion DNA polymerase (Thermo Fisher) and the amplicons obtained were introduced in pENTR207 or pENTR221 (Thermo Fisher) using Gateway BP Clonase II (Thermo Fisher). Transfer of inserts to destination vectors was done using Gateway LR Clonase II (Thermo Fisher). Point mutations were introduced using QuikChange® site-directed mutagenesis. All inserts were confirmed by DNA sequencing. The order of the different protein fusions (i.e. tag-REP or REP-tag) is indicated in the figures. Rep ORF from the TYLCV isolate ‘Almeria’ was used in this study, unless mentioned otherwise.

### Yeast two-hybrid assays

For the GAL4 Y2H assay, all gene fragments were cloned into Gateway-compatible variants of pGADT7 and pGBKT7 (Clontech) (50). Resulting plasmids were introduced into *Saccharomyces cerevisiae* strain PJ69-4α (51) using the standard lithium acetate/single-stranded DNA/polyethylene glycol 3,350 method (52). Transformed colonies were selected using minimal yeast medium (MM) lacking the amino acids Leu and Trp. To select for protein-protein interactions (PPIs), three independent transformants were randomly picked. After resuspension in 100 μl sterile double-distilled water, a 10-fold serial dilution was spotted on MM agar plates lacking Leu, Trp and His (*–*LWH) supplemented with 1 mM 3-Amino-1,2,4-triazole (3AT). The plates were incubated at 30°C for 3 days prior to scoring yeast growth.

The Gateway-ready plasmids for the split-ubiquitin Y2H system, pMET/pCub (53) were transformed into *S. cerevisiae* strain JD53. Selection of transformants and positive PPIs was performed as described (54). Transformed colonies were selected on yeast MM lacking His and Trp (–HW). To select for PPIs, the yeast colonies were resuspended, and a 10-fold serial dilution was spotted (similar to GAL4 assay) on MM agar plates supplemented with 0.1 mM CuSO_4_ and the appropriated amino acids. To increase selection specificity, 1 mg/mL 5-FOA (5-Fluoroorotic acid, Sigma) was added to the selection plates. Plates were incubated at 30°C for 4 days prior to scoring.

### Transient expression of proteins in N. benthamiana using agroinfiltration

All binary constructs were transformed in *Agrobacterium tumefaciens* (*Rhizobium radiobacter*) strain GV3101 (55) by electroporation (25 µF, 400 Ω, 1.25 kV/mm). Single colonies were grown overnight to OD600 0.8-1.5 in low salt LB broth (1% w/v Tryptone, 0.5% w/v yeast extract, 0.25% w/v NaCl, pH 7.0). Bacterial cells were pelleted by centrifugation (3000*g*) for 5 minutes, washed and resuspended in leaf infiltration medium (1× MS [Murashige and Skoog] salts, 10 mM MES pH 5.6, 2% w/v sucrose, 200 μM acetosyringone). *A. tumefaciens* carrying the pBIN61 binary vector to express the P19 silencing suppressor (referred to as pBIN61:P19) from Tomato busy shunt virus (TBSV) (56) was added to every experimental sample.

### Bimolecular fluorescence complementation (BiFC) assay

*Rep* from the TYLCV isolate ‘Alb13’ (*Rep*_Alb13_), the Arabidopsis *SUMO1* (AT4G26840) and *SCE1* (AT3G57870) coding sequence were cloned in the vectors pDEST-GWSCYCE to express fusions with the C-terminal half of S(CFP)3A (residues 156-239; referred to as SCFPC) or pDEST-SCYNEGW (N-terminal half of S(CFP)3A residues 1–173; SCFPN) (57). Four-week-old *N. benthamiana* leaves were syringe-infiltrated with *A. tumefaciens* suspensions at a final OD600 of 1.0 when a single construct was delivered. When two cultures were co-infiltrated for BiFC analysis, the cultures were mixed at a ratio 1:1 to a final OD600 of 1.0. *A. tumefaciens* carrying pBIN61:P19 was added to every experimental sample at a final OD600 of 0.5. Three days post-infiltration *N. benthamiana* leaf material was collected to analyze protein expression. SCFP fluorescence was detected using an excitation wavelength of 458 nm (argon laser), primary beam-splitting mirrors 458/514 nm, secondary beam splitter 515 nm and band filter BP 470-500 nm.

### In planta protein localization

*Rep*_Alb13_ was cloned in plasmid pGWB654 (C-terminal monomeric Red fluorescent protein (mRFP) tag) (58). The expression and detection were the same as for the BiFC assay.

### Confocal microscopy and image analysis

A confocal laser scanning microscope, Zeiss LSM510, was used to capture fluorescent images using a Zeiss c-Apochromat 40× 1.2 water-immersion Korr objective. Fluorescence was detected using the following beam and filter settings: GFP excitation at 488 nm (argon laser), primary beam-splitting mirrors 405/488, secondary beam splitter 490 nm, band filter BP 505-550 nm; RFP-excitation at 543 nm (helium-neon laser), primary beam-splitting mirrors 488/543 nm, and secondary beam splitter 635 nm, band filter LP 585-615 nm. For all observations, the pinhole was set at 1 Airy unit. The *Rep*_Alb13_ variant was used for the confocal work. For every experimental condition/treatment, at least three independent leaves were examined, and one representative image is shown (at least 50 nuclei were examined per sample). For each image analysis, absolute intensity values were used (Raw16.tif format). Images were analyzed and processed using ImageJ Fiji 1.0v (58, 59). Counting of the nuclear bodies was performed using pictures of 30 nuclei per treatment.

### *In planta* DNA replication activity assay using heterologous expression of Rep

To assess DNA replication by Rep, 3-5 weeks old *2IR-GFP N. benthamiana* (43) were co-infiltrated with *A. tumefaciens* to express *Rep-mRFP* variants inserted in pGWB654 (60) and *A. tumefaciens* carrying pBIN61:P19. *A. tumefaciens* cultures were mixed at a ratio of 2:1 to reach a final OD600 of 0.8 and 0.4, respectively, prior to infiltration. Four days post-agroinfiltration, GFP and RFP fluorescence were imaged using whole detached leaves mounted in a Chemidoc MP (BioRad) using the presettings “Alexa 488” and “Rhodamine”, respectively. To estimate the relative fluorescence intensity three leaves of the same age on different plants were agroinfiltrated and imaged. Mean relative fluorescence intensities (integrated density/area size) were calculated for each infiltrated area and the intensities were normalized per leaf by defining both the lowest fluorescence value (score 0, background signal) and the highest value (score 1.0). Visualization and ANOVA tests were done using Prism 9.0v (GraphPad).

### Quantification of extrachromosomal molecules using qPCR

To quantify the level of ECMs, leaf samples were snap-frozen in liquid nitrogen and stored at –80°C prior to processing. Frozen tissue was ground using a steel ball in a mill (TissueLyser II, Qiagen) and total DNA was extracted from approximately 30 mg of leaf tissue (*2IR-GFP N. benthamiana*) using the hexadecyltrimethylammonium bromide (CTAB) method (61). DNA concentrations were determined measuring the absorbance at 260nm on a Nanodrop (Thermo Fisher). In total, 250 ng of total DNA was used as template for a real time PCR reaction (QuantStudio3, Thermo Fisher) using the Hot FIREPol EvaGreen qPCR kit according to the suppliers’ instructions (Solis Biodyne). The ECM signal was normalized using the *25S rRNA* (Genbank ID, KP824745.1) as internal reference. Viral primers were designed such that they only amplify ECMs and not the *2IR-GFP* genomic insert (**Table 1**). All cycle threshold (Ct) values were corrected for primer efficiencies. All expression data was analyzed following the standard workflow provided by qBASE+ (Biogazelle) using three independent biological replicates (n=3). CNRQ was calculated using qBASE+. Data visualization and ANOVA tests were performed using Prism 9.0v (GraphPad).

**Table 1.**
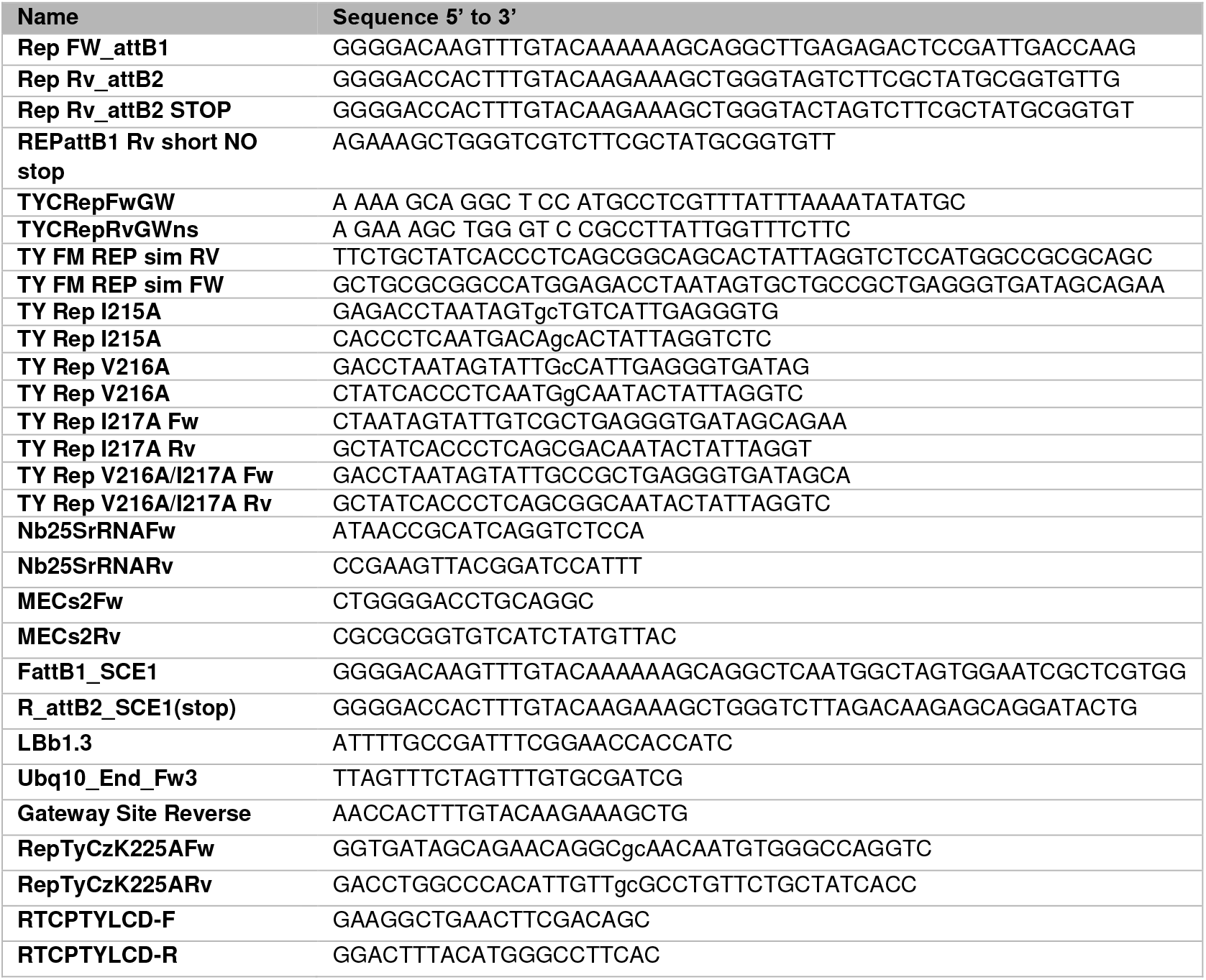
Sequences of the primers used in this study.

### Protein extraction from plant tissue and immunoblotting

Total protein fraction was extracted using approximately 30 mg of frozen *N. benthamiana* leaf tissue. Thereto, leaf material was snap frozen and grinded using plastic pestles after which RIPA buffer (50 mM Tris-HCl pH 7.4, 150 mM NaCl, 1% v/v Triton X-100, 0.5% w/v deoxycholate, 0.1% w/v SDS, 5 mM DTT, 1× cOmplete™ Protease Inhibitor Cocktail (Roche)) was added at a 4:1 v/w buffer/tissue ratio. Samples were thawed on ice and vortexed three times for 10 seconds. After incubating the homogenates prior on a rotating wheel for 1 hour (25 rpm) at 4°C, they were centrifugated at 16,000*g* (4°C) and 80 µL of the supernatant was mixed with 2× Laemmli buffer (100 mM Tris pH 6.8, 20% w/v glycerol, 4% w/v SDS, 100 mM DTT, 0.001% w/v Bromophenol blue) in 1:1 ratio. Total protein extract was denatured by heating at 96°C for 10 minutes. Upon protein denaturation, extracts were centrifugated at maximum speed (16,000*g*, ambient temperature) for 5 minutes and the supernatant was separated on a 12% SDS-PAGE gel and subsequently transferred onto a PVDF membrane (Immobilon-P, Millipore). Immunodetection of the proteins was performed according to standard protocols using the antibodies detailed (**Table 3)**. For detecting chemiluminescence (ECL), a home-made solution was used (0.1 M Tris-HCl pH 8.5, 1.25 mM luminol (Sigma-Aldrich) in DMSO, 0.2 mM *p*-Coumaric acid (Sigma-Aldrich) in DMSO, 0.01% v/v H_2_O_2_) and the signals were imaged using a Chemidoc MP (Bio-Rad) or a light-sensitive X-ray film (Fuji Super RX). Equal loading of the different protein extracts was confirmed by examining the Rubisco levels using Ponceau S staining of the membranes. The intensity of the Rubisco signal was also used to normalize the relative the GFP and mRFP protein levels.

### Quantification of viral infections

Infectious clones are described in detail in **Table 2**. *Agrobaterium* carrying the pGreen TY-LCV constructs were cultivated 24-30h to saturation (OD600 3.5-4) and then pelleted by centrifugation for 10 min (3000*g*) before being resuspended in fresh low salt LB medium without antibiotics to a final OD600 7.5-8. Then, they were introduced into 3-week-old *N. benthamiana* by injecting the suspension in an abaxial bud. Plants were maintained at 25°C with natural daylight supplemented to a 16h photoperiod with sodium lamps. Three weeks post inoculation, apical leaves were harvested near the apical shoot to isolate total DNA and quantify the relative viral accumulation of TYLCV coat protein in *N. benthamiana* by quantitative real time PCR. To confirm that TYLCV mutants did not revert back to wildtype clone by spontaneous mutations, the isolated virus samples were sequenced. Normalization for viral DNA using the ribosomal *RNA 25S* as internal reference. Primers used for real time quantification are detailed in **Table 1**. Real time PCR primers were designed to only amplify the circularized viral DNA.

**Table 2.**
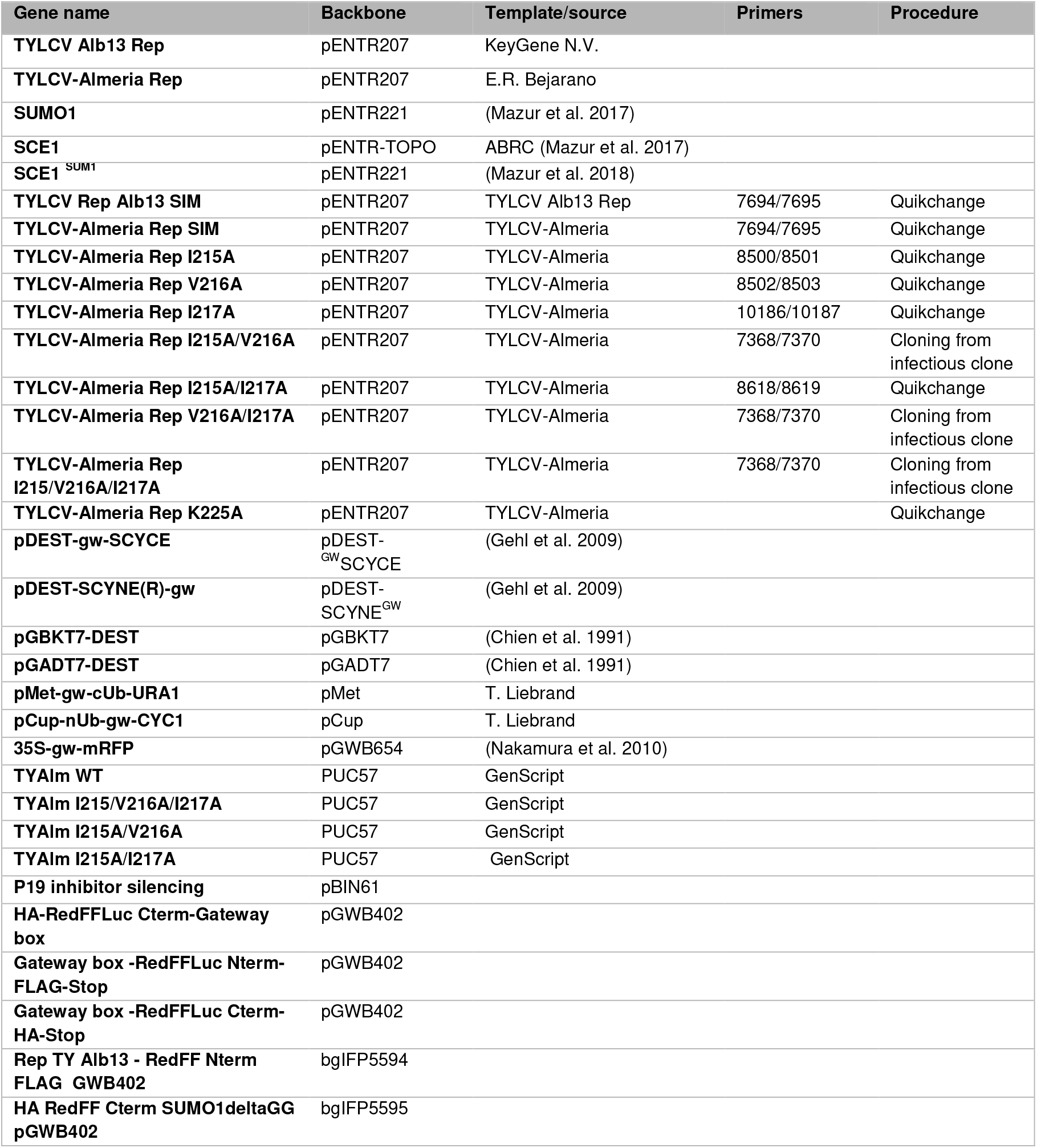
Plasmids used in this study.

**Table 3.** Antibodies used in this work.

| Antibody | Mono/polyclonal | Organism | Origin | Dilution |
| --- | --- | --- | --- | --- |
| **αGFP** | Monoc. | Rat | Chromotek [3H9] | 1:12000 |
| **αRFP** | Monoc. | Mouse | Chromotek [6G6] | 1:10000 |
| **αRat** | - | Goat | Pierce (31470) | 1:10000 |
| **αMouse** | - | Goat | Pierce (31430) | 1:10000 |

### Split-luciferase complementation (SLUC) assay

*N. benthamiana* leaves were co-infiltrated with a mixture of *A. tumefaciens* harbouring different pGWB402 (60) constructs (vectors used can be found in **Table 2**) and *A. tumefaciens* pBIN61:P19 with a final OD600 of 0.8 and 0.4 for both strains, respectively. *Rep*_Alb13_ variant was used for this SLUC. The infiltrated leaf areas were marked using a paint marker. Three days post-infiltration, agroinfiltrated leaves were brushed twice with D-luciferin buffer (235mM D-luciferin (Duchefa Biochemie) dissolved in Milli-Q water, 0.02% w/v Silwet L-77 (Crompton Europe). After 2 hours dark incubation, the chemiluminescence signal was captured using a CCD imaging system (Princeton Instruments) set at −70°C. An exposure time of 15 min with 10 × 10 binning was used for images. Data acquisition was performed using the MetaVue program (X-Rite). Each data point consisted of at least three replicates, and four independent experiments were performed for each assay. As negative control, unfused half of the luciferases were used. The log10 value of the mean signal (integrated density/area size) corrected for the leaf background signal and expressed as a proportion of total signal of all samples infiltrated in a single leaf was used for further statistical test. The data of each independent experiment were pooled for the ANOVA followed by Dunnett’s multiple comparison test in Prism v10.0 (GraphPad).

### Rep protein expression and purification

To express and purify recombinant Rep, the *E. coli* strain BL21-Gold (Agilent Technologies) was used harboring *Rep* in pGEX (N-terminal Glutathione S-transferase (GST)-tag, PreScission-cleavage site). Two-liter cultures were started with an overnight pre-culture to reach an OD600 of 0.1 and grown at 37°C, 200 rpm until reaching an OD600 of 0.7. Rep protein expression was induced using 0.5 mM IPTG and the temperature was lowered to 30°C. After two hours of induction, the bacterial cells were harvested at 5000*g* for 15 min at room temperature and the cell pellet was stored at −20°C prior to purification. The frozen pellet was thawed and resuspended in 30 mL of lysis buffer (50 mM Tris-HCl pH 7.5, 300 mM NaCl, 10% w/v glycerol, 2 mM MgCl_2_, 0.5 mM CaCl_2_, 2 mM DTT, 1% w/v Tween-20, 0.5 mg/mL lysozyme, 6 U/mL DNase I, and 1 mM PMSF) and stirred for 1 hour at 4°C. Cells were disrupted by passing them 3-5 times through a French press (Thermo Fisher) and centrifuged for 50,000*g* for one hour at 4°C. Supernatant was loaded on a 25 mL Glutathione Sepharose 4 Fast Flow column (Cytiva) and washed with 1.5 column volumes (CV) of wash buffer I (50 mM Tris-HCl pH 7.5, 200 mM NaCl, 5% w/v Glycerol, 2 mM MgCl2, and 2 mM DTT). Subsequently, the column was washed with 1.5 CV of wash buffer II (50 mM Tris-HCl pH 7.5, 800 mM NaCl, 5% w/v Glycerol, 2 mM MgCl_2_, and 2 mM DTT). The column was equilibrated with 1.5 CV of wash buffer I and 50 μL (8 mg/mL) of the His-tagged PreScission protease was added for overnight digestion. Protein elution was undertaken with 1.5 CV of elution buffer (50 mM Tris-HCl pH 7.5, 200 mM NaCl, 5% w/v glycerol, 2 mM MgCl_2_, 2 mM DTT, and 20 mM GSH). The eluted protein was further concentrated using a spin concentrator (10 kDa Amicon, Millipore) and rebuffered in 20 mM HEPES pH 7.5, 200 mM NaCl, 2 mM MgCl_2_, 5% w/v glycerol, and 2 mM DTT. To quantify the concentration of purified protein in the samples a Bicinchoninic acid (BCA) assay was conducted using the BCA kit following the manufacturer recommendations (Sigma Aldrich).

### ATPase activity assay

The decrease in NADH absorbance, which is proportional to the rate of ATP hydrolysis, was measured continuously at a wavelength of 340 nm for 120 minutes at 20°C with the Synergy H1MF (BioTek) plate reader. The assay was performed in 25 mM HEPES/NaOH pH 7.5, 200 mM NaCl, 10 mM KCl, 10 mM MgCl_2_, 1 mM DTT, 5% w/v glycerol, 1.5 mM NADH, 10 U each of pyruvate kinase (PK) and lactate dehydrogenase (LDH) and 3 mM of phosphoenolpyruvate (PEP). The quantity of sample was adjusted to achieve complete consumption of NADH within the measurement time. For wildtype Rep, Rep I217A, Rep I215/217A, Rep V216/I217A, Rep K225A, we used a final concentration of 8 μM, 80 μM, 80 μM, 130 μM and 50 μM of protein, respectively, to observe ATP hydrolysis within the measurement time. Experimental replicates were measured. The negative control contained all the reaction components except for the Rep protein. To each well, 1 mM ATP was added to start ATP hydrolysis.

### Determination of ATP turnover rate

A linear decrease of NADH from each measurement (described above), spanning over at least 15 min (62), was selected to calculate decrease in absorbance over time, *i.e.* the slope of the curve. An NADH standard curve ranging from 0.0 mM to 2.0 mM NADH was used to correlate the absorbance values at 340 nm with the respective NADH concentrations. Hence, the slope of the curve (absorbance decrease per time interval) could be transformed into NADH consumption over time (mM/min). Spontaneous oxidation of NADH observed in the negative control, i.e. a mock sample without Rep, was subtracted from the corresponding absorbance values. The slope, intercept and R-squared values were determined by fitting a linear model using RStudio (R version 4.1.0.)

### Size exclusion chromatography

0.6-2.5 mg of Rep (2.5 mg WT Rep, 1.1 mg I217A, 1.9 mg I215/217A, 0.6 mg V216/I217A, 0.6 mg K225A) were diluted to a final volume of 100 µl each in 20 mM HEPES/NaOH pH 7.5, 200 mM NaCl, 2 mM MgCl_2_, 2.5% w/v glycerol, 2 mM DTT. The samples were loaded onto a Superdex 200 Increase 10/300 GL (Cytiva) column that had previously been equilibrated with the same buffer and was attached to an Äkta^TM^ pure FPLC system (Cytiva). Isocratic elution was performed over 1.25 column volumes and fractions of 0.5 mL were collected across the entire width of each peak. The fractions were pooled regarding their allocation to a certain (sub-)peak and used for further analysis, including SDS-PAGE and tandem mass spectrometry.

### Accession numbers

DNA clones of TYLCV isolate ‘Alb13’ *Rep* (Genebank ID: FJ956702.1) were kindly provided by Keygene (Wageningen, Netherlands); *Rep* from TYLCV isolate ‘Almeria’ ( AJ489258), CtLCV, Cotton leaf curl virus *Rep* (KC412251.1), BGMV, Bean golden mosaic virus *Rep* (JF694454.1); CaLCV, Cabbage leaf curl virus *Rep* (U65529.2), PepGMV, Pepper golden mosaic virus *Rep* (EF210556), SLCCNV, Squash leaf curl China virus *Rep* (KC222956.1) and the TYLCV infectious clone (based on AJ489258.1) were synthetized by GenScript. Clones for the coding sequences of *AtSCE1* (At3g57870) and *AtSUMO1* (At4g26840) were previously described (71). For the Rep protein alignment, we included the Rep protein sequences from TGMV, Tomato golden mosaic virus (NC001507); ACMV, African cassava mosaic virus (CAD20827.1) and BCTV, Beet curly top virus (AAK59260.1).

## Acknowledgements

We are grateful to L. Tikovsky and H. Lemereis for excellent plant care; LCAM-FNWI (University of Amsterdam) for maintenance and help with confocal microscopy. We thank T. Nagakawa (Shimane University, Japan), J. Kudla (WWU Munster, Germany) for sharing plasmids. We thank L. Deslandes (LIPME Toulouse, France) for sharing the yeast strain JD53. This work was supported by the Spanish Ministerio de Ciencia y Tecnología (PID2022-139376OB-C31) (AP-L, BS and ER-B), the Topsector T&U grants 1409-036 and LWV19157 to HvdB including the partnering breeding companies.

## Supplementary

**Supplementary figure 1:**
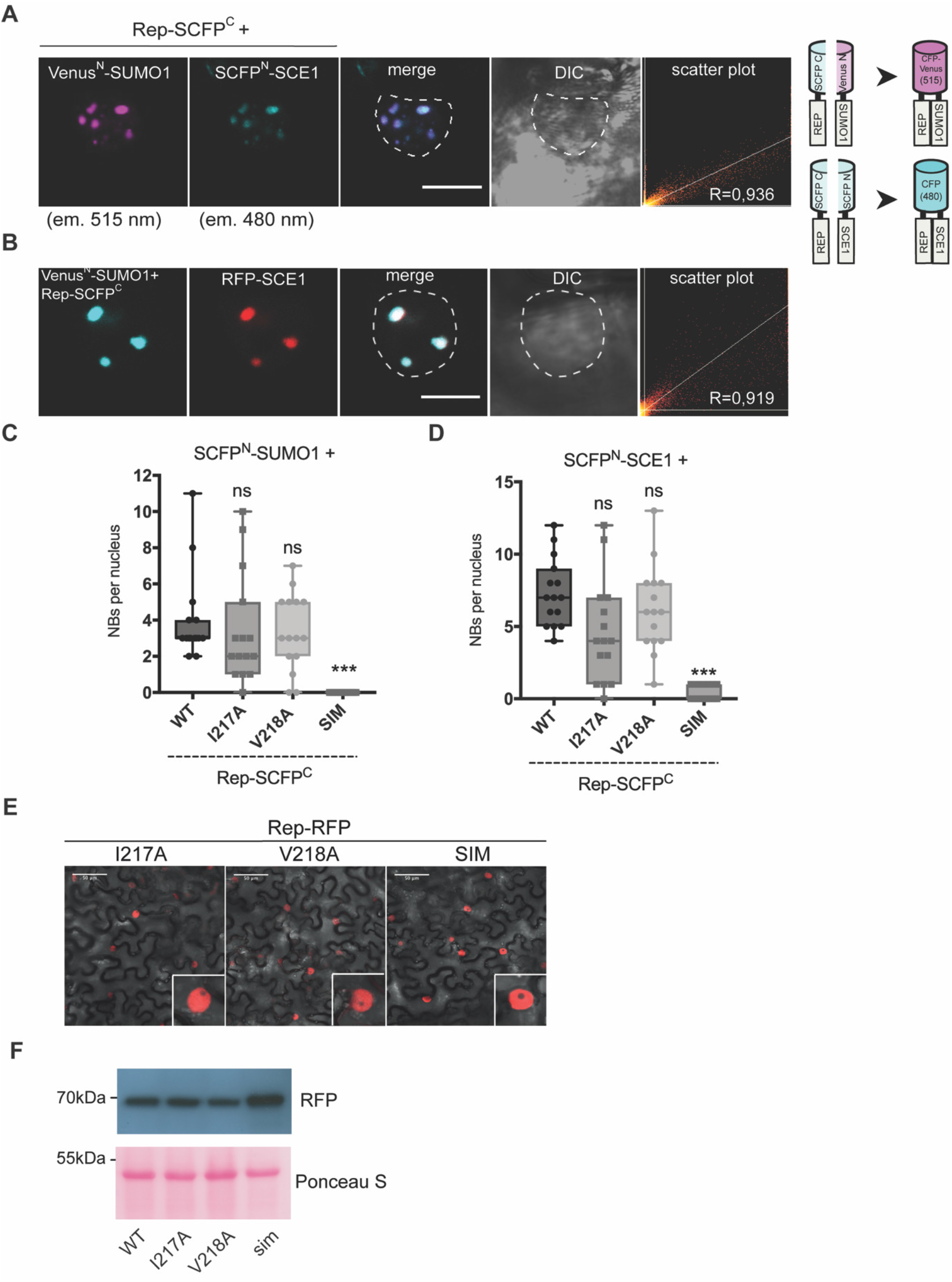
Rep interacts dynamically with SUM0 and SCE1 and contains a conserved SUM0 Interacting Motif -.

**Supplementary figure 2:**
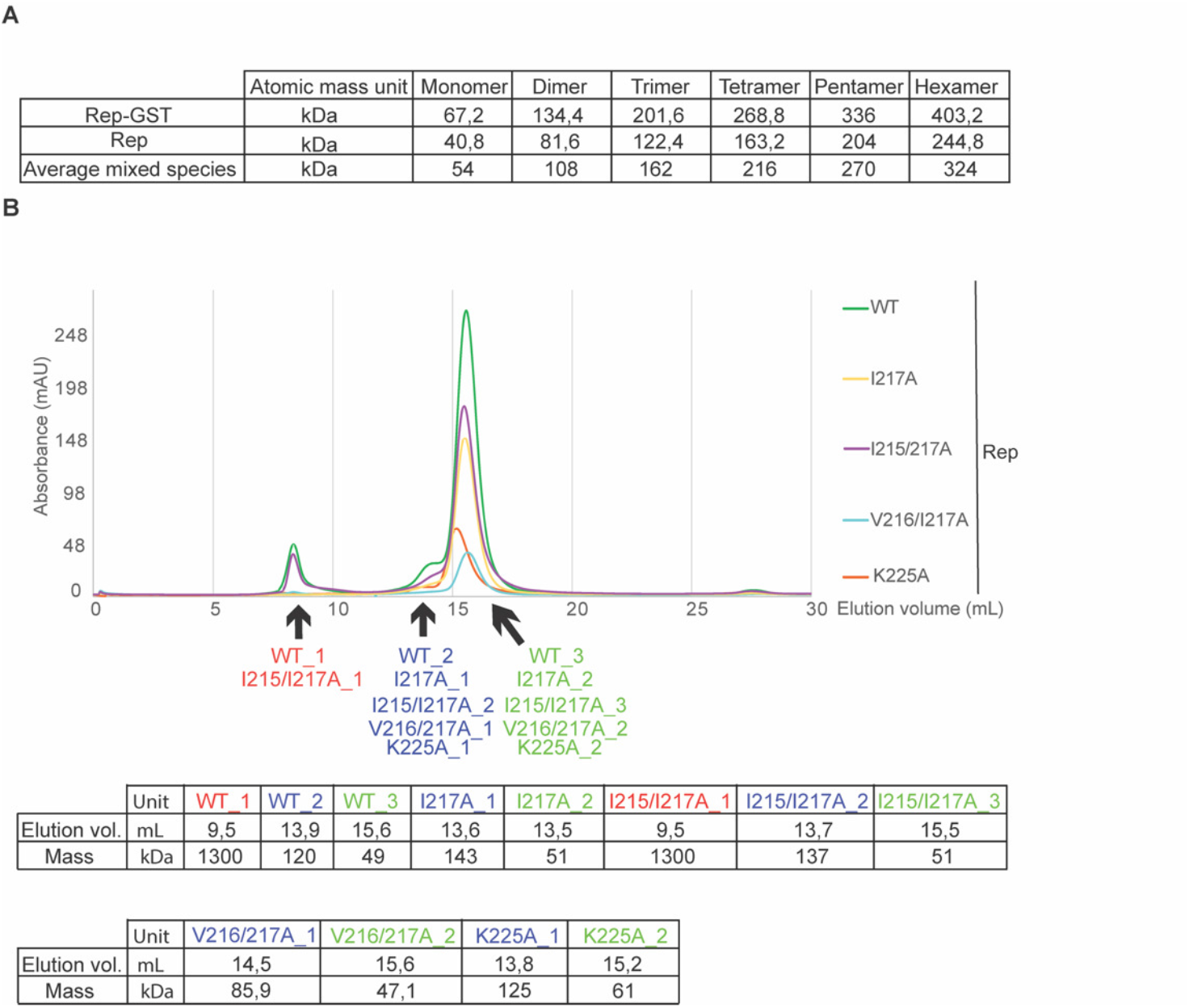
Detailed analytical size exclusion chromatography of Rep SIM variants.

